# Subcellular protein localisation of *Trypanosoma brucei* bloodstream form-upregulated proteins maps stage-specific adaptations

**DOI:** 10.1101/2022.11.07.515401

**Authors:** Clare Halliday, Samuel Dean, Jack Daniel Sunter, Richard John Wheeler

## Abstract

Genome-wide subcellular protein localisation in *Trypanosoma brucei*, through our TrypTag project, has comprehensively dissected the molecular organisation of this important pathogen. Powerful as this resource is, *T. brucei* has multiple developmental forms and we previously only analysed the procyclic form. This is an insect life cycle stage, leaving the mammalian bloodstream form unanalysed. The expectation is that between life stages protein localisation would not change dramatically (completely unchanged or shifting to analogous stage-specific structures). However, this has not been specifically tested. Similarly, which organelles tend to contain proteins with stage-specific expression can be predicted from known stage specific adaptations but has not been comprehensively tested. Here, we used endogenous tagging to determine the sub-cellular localisation of the majority of proteins encoded by transcripts significantly upregulated in the bloodstream form and compared them to the existing localisation data in procyclic forms. This confirmed the localisation of known and identified the localisation of novel stage-specific proteins, giving a map of which organelles tend to contain stage specific proteins: the mitochondrion for the procyclic form, and the endoplasmic reticulum, endocytic system and cell surface in the bloodstream form.

## Introduction

*Trypanosoma brucei* is a unicellular eukaryotic parasite and, like any unicellular organism, adjusts its gene expression profile to adapt to different environments. As an obligate parasite, the environments it encounters are exclusively within the host and vector and gene expression profile changes give rise to the appropriate protein machinery to adapt the parasite to these niches. *T. brucei* has three main replicative life cycle stages: the procyclic form (PCF, fly midgut), the epimastigote form (EMF, fly salivary glands) and the bloodstream form (BSF, mammalian host bloodstream), although within these stages there is also additional specialisation [1,2]. The PCF and BSF are readily grown in culture.

The PCF and BSF have many well characterised differences, including the BSF VSG surface coat and associated expression machinery [3], metabolic differences and associated remodelling of the mitochondrion [4], morphology, and morphogenesis adaptations [5,6], along with many more. However, genome-wide mapping of the global changes are broadly limited to gene expression level, most extensively determined at the mRNA level [7–11] which does not correlate fully with protein abundance [12]. Few studies consider later steps in protein production: translation (mRNA ribosome footprinting) [13] and protein abundance (quantitative proteomics) [14]. Despite the comparative ease of culturing PCFs and BSFs and the powerful reverse genetic tools available, a huge number of genes with evidence for BSF upregulation are not characterised.

Here, we aim to address this using subcellular protein localisations. We have demonstrated the power of this approach in PCFs with the TrypTag genome-wide protein localisation project [15]. This showed how informative localisation can be for holistic mapping of potential protein function, although naturally does not determine specific molecular function. We also previously used high throughput tagging BSF-upregulated genes to identify ESB1, necessary for transcription of the expression site containing the VSG gene along with expression site associated genes [16]. However, our previous analysis of these BSF localisations was minimal, aiming only to identify expression site body components. Here, we present analysis of an extended version of this BSF localisation dataset as both evidence for how BSFs are adapted relative to PCFs and as a resource for the research community.

## Results and Discussion

To enrich for proteins likely to have BSF-specific functions, we devised three gene tagging sets based on data available at the time (Figure 1). Set 1) 289 genes with mRNAs upregulated in BSFs, based primarily on mRNAseq from [7] but manually incorporating some genes identified as strongly upregulated in [8–11] not in [7]. Set 2) *T. brucei-*specific genes (defined as those which lack both an *L. major* Friedlin and *T. cruzi* Brener non-Esmareldo ortholog) not already included in Set 1, which met one of two criteria based on TrypTag PCF tagging data available at the time: Set 2a) the 30 genes that had failed to give a convincing signal above background by both N and C terminal tagging, and Set 2b) the 21 genes which had a nucleoplasm or nucleolar localisation. The former were selected to test whether lack of PCF signal correlated with BSF stage-specific expression, and the latter as candidates for *T. brucei-*specific BSF nuclear structure adaptation.

**Figure 1.**
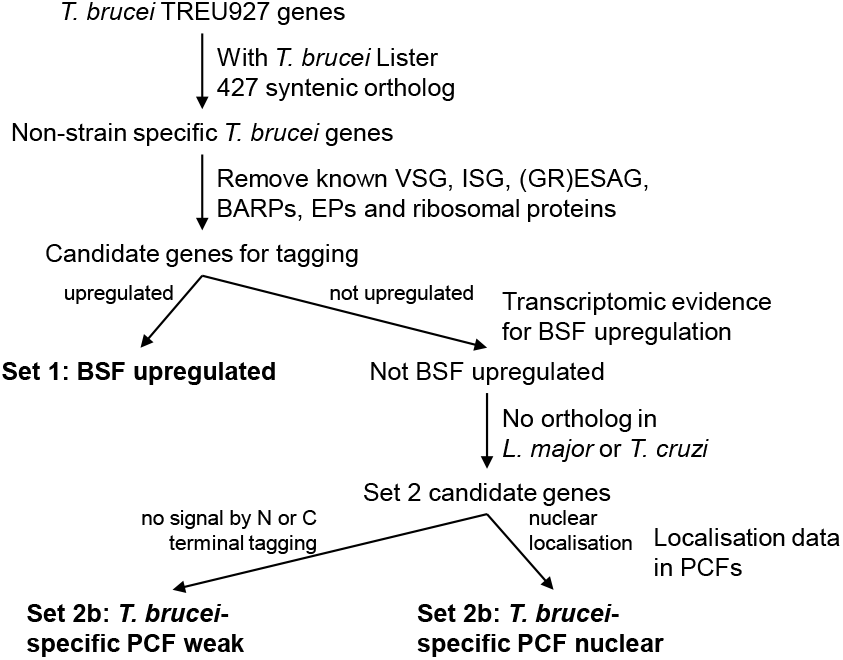
Flowchart for selection of genes for tagging in BSFs.

We prioritised N terminal tagging because this preserves the 3’ untranslated region (UTR), suspected to confer most gene regulation in trypanosomes [17]. However, when a protein had a predicted N terminal signal peptide C terminal tagging was instead necessary. If we failed to generate a drug resistant population, we repeated construct generation and transfection at least once. The final success rate generating cell lines (listed in Table S1) was 72.9% (Figure 2A), of which 76.6% had signal we manually classified as unlike background fluorescent signal (Figure 2B) – i.e. a convincing subcellular localisation.

**Figure 2.**
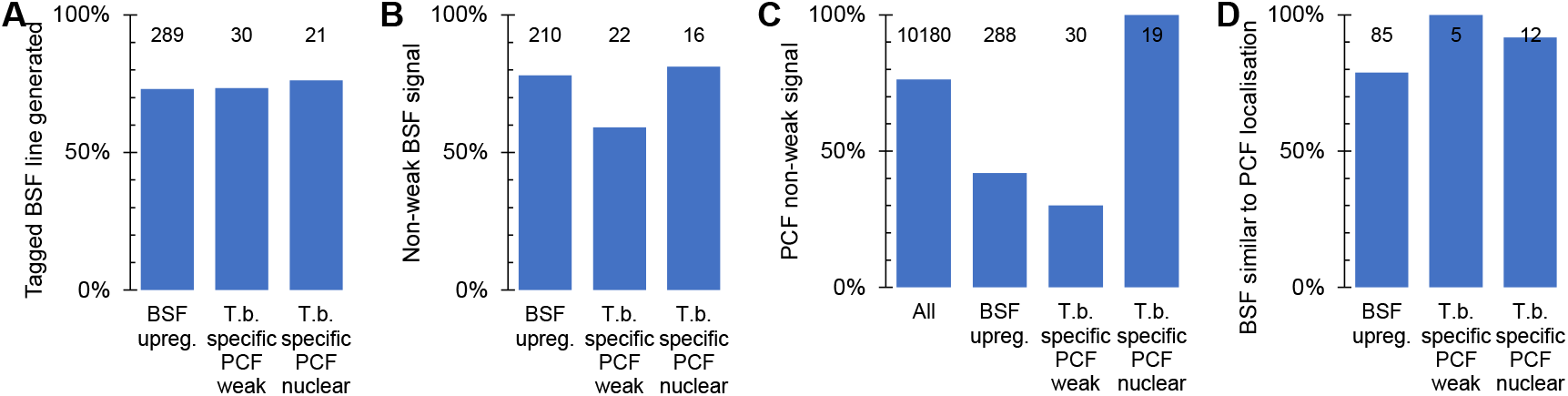
Success rates generating BSF localisations. **A**. Proportion of cell lines generated for each target gene set: Set 1, BSF upregulated; set 2a, *T. brucei*-specific with PCF weak signal by N and C terminal tagging; set 2b, *T. brucei*-specific proteins which localise to the nucleus in PCFs. **B**. Proportion of BSF cell lines generated for each target gene set which had a weak signal, i.e. no strong localisation to an identifiable organelle in the BSF. **C**. Proportion of each target gene set for which strong localisation to an identifiable organelle was observed for either N or C terminal tagging in PCFs, in comparison to all genes with PCF data. Data from the TrypTag project. **D**. The proportion of BSF cell lines for each target gene set with strong localisation to an identifiable organelle which gave a similar localisation to either N or C terminal tagging in PCFs. In each graph, the number of genes for which data is available in each group is shown at the top of each column.

For the final analysis of these localisations, we re-analysed the gene sets based on the entire TrypTag PCF localisation dataset [15] and TriTrypDB version 59 [18] (Figure S1). There were some changes; altered OrthoMCL sensitivity due to addition of new genomes (Figure S1B), additional PCF tagging repeats providing a strong convincing localisation where only weak signal was previously observed (Figure S1C), and changed PCF localisation annotation (e.g. from nucleoplasm to nuclear envelope, Figure S1D). We also defined a final criterion for upregulation in the BSF: transcripts significantly upregulated (*p* < 0.05, Student’s T test) by mRNAseq in the BSF relative to the PCF (data from [7]). However, overall, the gene sets well reflect their original purpose.

We observed convincing fluorescent signal in BSFs for many tagged proteins in Set 1 (upregulated in BSFs at the mRNA level, Figure 2B). In this gene set, disproportionately many genes were also *T. brucei*-specific (Figure S1B), with no convincing above-background localisation observed in the PCF (Figure S1C). We also observed a convincing fluorescent signal in BSFs for many in Set 2a (*T. brucei*-specific genes with no detectable PCF signal, Figure 2B). Lack of fluorescent signal in the PCF tagging previously raised our suspicions that these genes may not be expressed in this life cycle stage, never expressed, or encode a non-functional, and therefore degraded, protein product. Similarly, failure to generate a PCF tagged cell line may indicate inaccurate sequence data for that locus or that the drug selectable marker cannot be expressed from that locus. This was an acute concern when the gene was *T. brucei* specific and therefore had no evidence from evolutionary conservation for being functional. Our BSF localisation provides evidence that many of these genes encode an expressed and likely functional protein (on the basis that the proteins often targeted to a specific organelle), supporting proteomic analyses [19]. As would be expected, fluorescent signal in a tagged cell line therefore broadly correlates with mRNA abundance across life cycle stages and failure to observe a convincing localisation in PCFs is, as we previously proposed [15], at least partially predictive of a stage specific protein expression.

With the exception of Set 2b, the set of *T. brucei* specific nuclear genes which were selected based on a specific PCF localisation, our BSF tagging was of proteins disproportionately more likely to have no detected signal from PCF tagging (Figure 2C). However, when a PCF localisation was available it was likely to be similar to the BSF localisation we observed, overall ∼85% were manually classified as similar (Figure 2D). When dissimilar, the localisation observed in either the PCF or BSF was typically either a weak cytoplasmic signal or a cytoplasm, nuclear lumen and flagellar cytoplasm localisation. The former is simply background signal. The latter is the localisation we observed in PCFs for mNG when not fused to a protein, perhaps indicative that these dissimilarities are technical tagging failures or poorly tolerated fusion proteins – truncated or partially degraded leading to expression of effectively mNG alone. One, however, featured a clear change; see below.

For Set 1, the set of BSF upregulated genes, whether or not a PCF localisation was visible the BSF localisation gave a much stronger signal – detectable as we used the same microscope, camera and image processing settings for PCFs and BSFs, making signal intensity in the images approximately quantitative. This includes proteins known or expected to be BSF-upregulated: pyruvate transporter 1, PT1 [20]; repressor of differentiation kinase 2, RDK2 [21]; flagellum adhesion protein 3, FLA3 [22]; and cytoskeleton associated protein CAP5.5V [23] (Figure 3A). However, it also includes many novel or uncharacterised proteins localising to many different organelles (examples in Figure 3B). We also noted one clear example where protein localisation differed between the PCF and BSF. Tb927.11.1230 and its syntenic ortholog Tb427tmp.47.0026 localised to the distal axoneme (occasionally with weak proximal signal) in PCFs and the entire axoneme in BSFs. Interestingly, this is comparable to the flagellar protein FLAM8 (flagellar member 8) [24].

**Figure 3.**
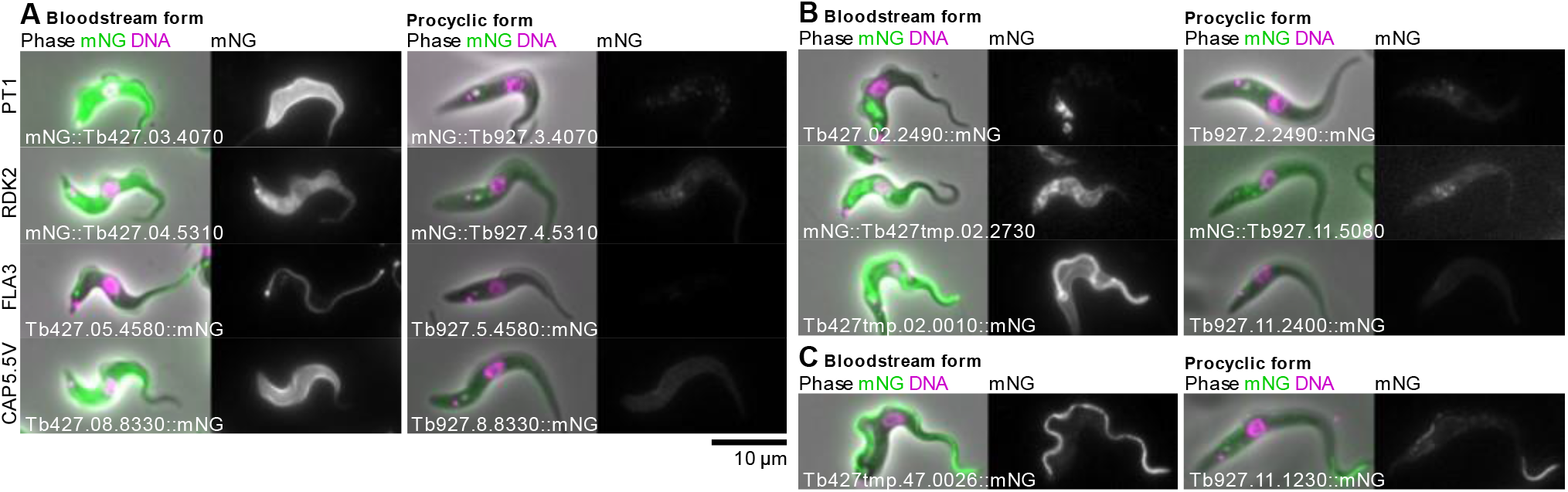
Example subcellular localisations of BSF-specific or strongly upregulated proteins. **A**. Known or expected BSF-specific proteins, showing the BSF localisation in *T. brucei* Lister 427 on the left and the localisation of the *T. brucei* TREU927 ortholog in PCFs from the TrypTag project on the right. For each cell line, an overlay of the phase contrast, mNG fluorescence and the Hoechst 33342 DNA stain is shown on the left and the mNG fluorescence alone in greyscale on the right. The gene ID and mNG fusion is shown in the bottom left. BSF and PCF mNG fluorescence are shown at approximately equal contrast levels to enable comparison of protein levels. **B**. Examples of previously uncharacterised BSF-specific proteins localising to (from top to bottom) the endocytic system, the endoplasmic reticulum and the pellicular and flagellar membranes. **C**. Only identified example of a protein whose subcellular localisation differs between BSFs and PCFs, localising to the whole axoneme in BSFs concentrated in the distal axoneme in PCFs.

We noted that BSF-upregulated proteins often localised to membranous structures - the pellicular or flagellar membrane, the endoplasmic reticulum or the endocytic system (Figure 3A,B). We therefore tested for a bias in localisation annotation term usage relative to genome-wide usage in PCFs. Taking only the target genes for BSF tagging not selected based on a nuclear PCF localisation, i.e. excluding Set 2b, there was indeed a significant bias in term usage (*p* < 10^−30^). Normalised fold-change in usage of annotation terms revealed a strongly disproportionately high usage of terms associated with the surface membrane and the endo/exocytic system (pellicular and flagellar membrane, ER and endocytic). There were also weaker biases in BSFs for 1) general (nucleus, nuclear lumen) rather than specific (nucleoplasm, nucleolus) nuclear localisation annotations, 2) fewer mitochondrion and kinetoplast annotations, 3) more glycosome terms, and 4) more flagellum tip and flagellar connector-like [5,25] annotation terms (Figure 4A,B). The BSF cell surface therefore has the greatest adaptation between BSFs and PCFs, with this change plausibly supported and/or maintained by changes in the ER and endocytic system.

**Figure 4.**
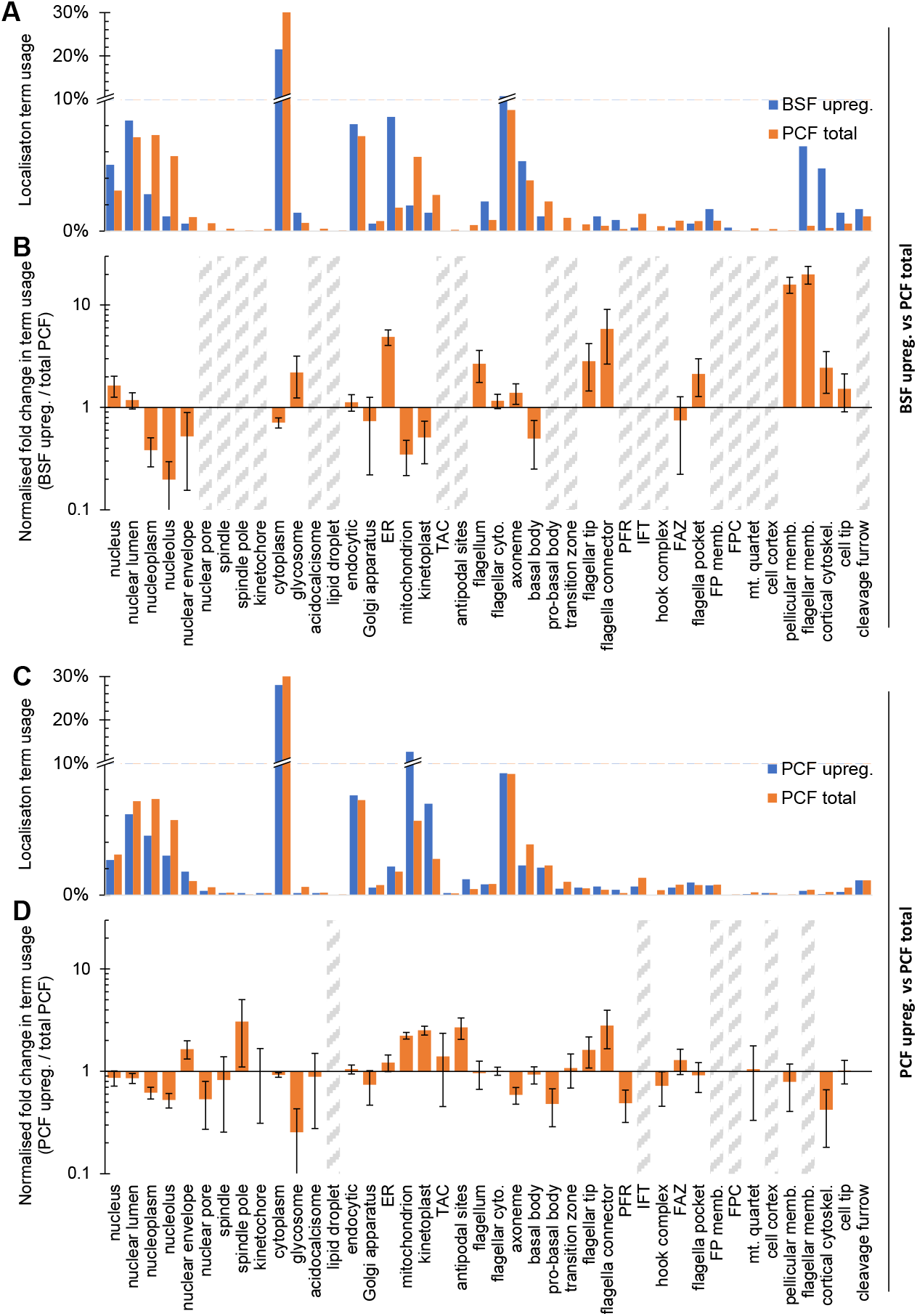
Stage-specific organelle adaptation mapped using localisation term usage. **A**. Localisation annotation term usage, as the proportion of all annotation terms used localisations, comparing all PCF (N and C terminal tagging) localisation terms to all BSF localisations described here, excluding the target gene set 2b; *T. brucei*-specific nuclear localising proteins. All localisation annotations for N and/or C terminal tagging, whichever are available, so long as they did not have the ‘weak’ or ‘<10%’ modifiers. **B**. The data in **A**, except plotted as the ratio of term usage in BSF upregulated *vs*. total PCF, normalised to number of annotation terms in the BSF set. Error bars represent standard error of proportion. Grey hatched bars indicate too few (<3) BSF upregulated protein localisations for accurate fold change calculation. **C**. Analogous analysis of PCF upregulated genes from TrypTag data: Localisation annotation term usage, as the proportion of all annotation terms used for non-weak localisation, comparing all PCF localisation terms with those for proteins encoded by genes significantly upregulated at the mRNA level in PCFs. **D**. The data in **C**, except plotted as the ratio of term usage in PCF upregulated *vs*. PCF total term usage, normalised to number of terms in the PCF set. Error bars represent standard error of proportion. Grey hatched bars indicate too few (<3) PCF upregulated protein localisations for accurate fold change calculation.

The converse analysis, taking genes upregulated in the procyclic form [7] and analysing localisation annotation term usage relative to genome-wide usage in PCFs also revealed a significant change (*p* < 10^−30^) in term usage, reflecting adaptation in the PCF. We identified 1) disproportionately high usage of mitochondrion and kinetoplast terms, 2) high usage of flagellar tip and flagellar connector terms, and 3) few glycosome terms. This speaks to the known upregulation of oxidative phosphorylation (mitochondrial) relative to glycolysis (glycosomal) as the major ATP source in procyclic form and adaptation of the flagellum tip likely linked with new flagellum outgrowth [5], but limited other changes (Figure 4C,D).

In conclusion, we have mapped which organelles contain proteins upregulated in the *T. brucei* BSF and PCF life cycle stages (summarised in Figure 5), thus mapping where the molecular machinery responsible for their stage-specific adaptations likely act in the cell. This includes many uncharacterised proteins with little or no bioinformatic insight into likely function. Lack of fluorescent signal by endogenous tagging in the PCF was often predictive of BSF expression, confirming the power of the TrypTag genome-wide protein localisation resource as a protein expression level resource. We also showed that it is likely that a large majority of *T. brucei* proteins, when expressed, have similar localisations in BSFs and PCFs – the dominant adaptive process therefore appears to be change in expression level rather than change in localisation. We suggest that this also likely applies to other life cycle stages and the different life cycle stages of other trypanosomatid parasites.

**Figure 5.**
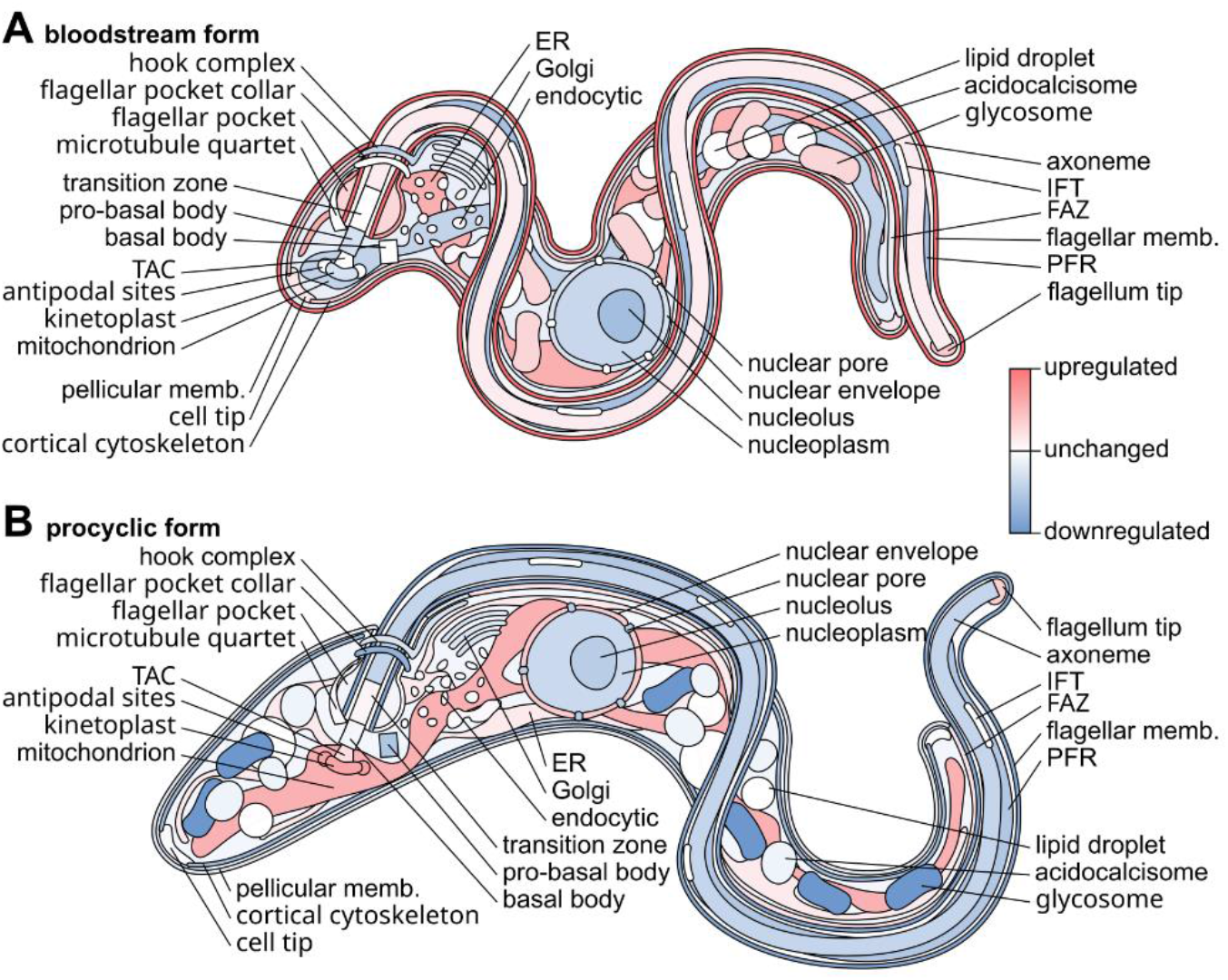
Diagrammatic summary of stage-specific organellar adaptation. Diagrammatic representation of G1 *T. brucei* cell structure with organelles colour-coded by whether they contain disproportionately many genes with stage-specific expression level. **A**. Summary of organelles tending to contain proteins with BSF-specific up or downregulation, data from Figure 4B. **B**. Summary of organelles tending to contain proteins with PCF-specific up or downregulation, data from Figure 4C.

## Methods

### Cell culture

Bloodstream form *Trypanosoma brucei brucei* strain Lister 427 pJ1339 was grown in HMI-9 at 37°C with 5% CO_2_ [26], maintained in log phase growth and at less than ∼2×10^6^ cells/ml by regular subculture. To enable CRISPR/Cas9 genome modifications, this cell line expresses T7 RNA polymerase, Tet repressor, Cas9 nuclease and puromycin drug selectable marker [16] and were maintained with periodic drug selection using 0.2 µg/ml Puromycin Dihydrochloride. Culture density was measured with a CASY model TT cell counter (Roche Diagnostics) with a 60 µm capillary and exclusion of particles with a pseudo diameter below 2.0 µm.

### Electroporation and drug selection

For endogenous tagging of a protein, electroporation was used to transfect *T. brucei* with two linear DNA constructs; one from which a CRISPR sgRNA is transiently expressed and one carrying the fluorescent protein and drug selectable marker which has homology arms allowing homologous recombination into the target locus. Constructs for endogenous N or C terminal tagging constructs were generated using long primer PCR from a pPOTv7 mNeonGreen (mNG) / blasticidin deaminase template, and PCR was used to generate DNA encoding sgRNA with a T7 promoter, both as previously described [27,28] (primer sequences in Table S1). ∼5 µg of DNA from the PCRs was purified by phenol chloroform extraction, resuspended in 10 µl water, then mixed with approximately 3×10^7^ cells resuspended in 100 µl of Roditi Tb-BSF buffer [29]. Transfection was carried out using program X-001 of the Amaxa Nucleofector IIb (Lonza) electroporator in 2 mm gap cuvettes. Following electroporation, cells were transferred to 10 ml pre-warmed HMI-9 for 6 h then 5.0 µg/ml Blasticidin S Hydrochloride added to select for cells with successful construct integration. Healthy resulting populations were maintained with periodic drug selection using 0.2 µg/ml Puromycin Dihydrochloride and 5.0 µg/ml Blasticidin S Hydrochloride.

### Selection of genes for tagging

BSF tagging was carried out in the *T. brucei* Lister 427 cell line, and we considered genes for tagging if they had a syntenic ortholog in *T. brucei* TREU927. Genes were selected for tagging as described in the main text using TrypTag PCF protein localisation data available up to 12^th^ March 2018 and TriTrypDB version 36, with the following specific exclusion criteria to avoid tagging of large well-known gene families and genes encoding GPI-anchored proteins known to be refractory to N and C terminal tagging. VSG, the major BSF surface coat protein was excluded by removing known (named) VSG genes and pseudogenes. In the interest of unbiased analysis, we ensured surface coat proteins characteristic of other life cycle stages were also excluded: EP procyclins, also called procyclic acidic repetitive proteins (PARPs), and *brucei* alanine rich proteins (BARPs). Known (named) invariant surface glycoproteins (ISGs) were excluded, with the exception of tagging controls ISG65 and GPI-PLC, and VSG expression site associated genes and related genes (ESAGs and GRESAGs) were excluded. Finally ribosomal proteins, which we deemed unlikely to be of interest, were excluded.

### Light microscopy

Cells were prepared for light microscopy by centrifugation to remove medium, followed by resuspension in FCS-free HMI-9 containing 1 µg/mL Hoechst 33342 before a second centrifugation and resuspension in a small volume (∼20 µl) of FCS-free HMI-9. An equal volume of 0.04% (v/v) formaldehyde in FCS free HMI-9 was added to lightly fix the cells [16,30]. Images were captured on a DM5500 B (Leica) upright widefield epifluorescence microscope using a plan apo NA/1.4 63× phase contrast oil-immersion objective (Leica, 15506351) and a Neo v5.5 (Andor) sCMOS camera using MicroManager [31].

### Statistics

Statistical significance of change in localisation annotation terms usage for the PCF and BSF upregulated gene sets was evaluated using the Chi squared test, taking the annotation term usage in the genome-wide PCF set as the null hypothesis. Fold change in individual term usage was calculated as the ratio of term count in the PCF or BSF upregulated set to the term count in the genome-wide PCF set, eg. count of axoneme annotation terms in the BSF upregulated gene set divided by count of axoneme annotation terms genome-wide in PCFs. This is an approximation for BSFs, as we do not know the genome-wide term usage in BSFs. Error was estimated using the standard error of proportion (SEP) for each annotation term. Fold change in term usage was normalised (and SEP scaled appropriately) to the total number of annotation terms in each set, such that no bias in usage between sets is unity.

## Acknowledgements

We would like to thank the Wellcome Trust for supporting this work through a Sir Henry Dale Fellowship to RJW [211075/Z/18/Z] and the biomedical resources grant which supported TrypTag [108445/Z/15/Z]. We would like to thank Keith Gull and the TrypTag principal investigators team for their support in TrypTag-associated projects.

## Data availability

All microscopy data is available at Zenodo indexed under DOI: 10.5281/zenodo.7258722 [32].

## Supplemental

Table S1. **Localisations and primer sequences**.

**Figure S1.**
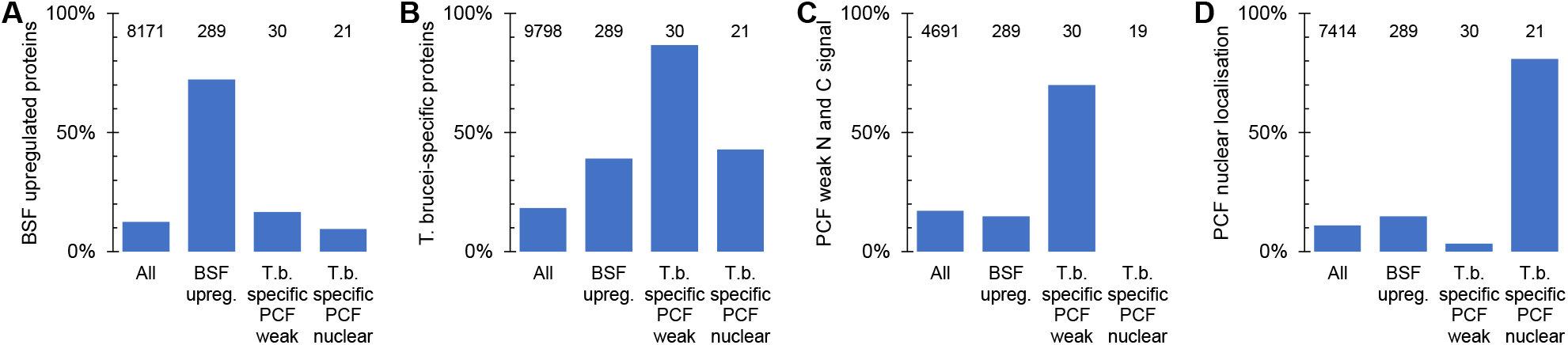
*Post hoc* analysis of the target gene sets for BSF tagging. **A**. Proportion of genes at least 2.5-fold upregulated mRNA and *p* < 0.05 (two-tailed T test) from [7], for each target gene set; BSF upregulated, *T. brucei*-specific with PCF weak signal by N and C terminal tagging and *T. brucei*-specific proteins which localise to the nucleus in PCFs, in comparison to all *T. brucei* genes. **B**. Proportion of genes with no *L. major* and no *T. cruzi* ortholog in each target gene set. **C**. Proportion of genes with N and C terminal tagging data in PCFs from the TrypTag project for which both termini had weak, i.e. no strong localisation to an identifiable organelle. **D**. Proportion of genes annotated as localising to the nucleus, nucleoplasm or nucleolus by either N or C terminal tagging in PCFs from the TrypTag project. In each graph, the number of genes for which data is available in each group is shown at the top of each column.

